# Next-generation matrices for marine metapopulations: the case of sea lice on salmon farms^*^

**DOI:** 10.1101/2022.04.25.489382

**Authors:** Peter D. Harrington, Danielle L. Cantrell, Mark A. Lewis

## Abstract

Classifying habitat patches as sources or sinks and determining metapopulation persistence requires coupling connectivity between habitat patches with local demographic rates. In this paper we show how next-generation matrices, originally popularized in epidemiology to calculate new infections after one generation, can be used in an ecological context to couple connectivity with local demography to calculate sources and sinks as well as metapopulation persistence in marine metapopulations. To demonstrate the utility of the method, we construct a next-generation matrix for a network of sea lice populations on salmon farms in the Broughton Archipelago, BC, an intensive salmon farming region on the west coast of Canada where certain salmon farms are currently being removed under an agreement between local First Nations and the provincial government. We identify the salmon farms which are acting as the largest sources of sea lice and show that in this region the most productive sea lice populations are also the most connected. We find that the farms which are the largest sources of sea lice have not yet been removed from the Broughton Archipelago, and that warming temperatures could lead to increased sea louse growth.

## 1 Introduction

Metapopulations consist of subpopulations located on isolated habitat patches that are connected via dispersal [1, 2, 3]. In most benthic marine species, this dispersal comes from the pelagic larval stage [4]. Larvae disperse between, and then settle on habitat patches, and once settled the remaining stages are sedentary and remain confined to a specific habitat patch. In marine systems the metapopulation concept, where subpopulations are connected but have their own demographic rates, has been used in the spatial planning of Marine Protected Areas and the siting of marine reserves [5, 6, 7, 8, 9, 10].

In a metapopulation framework, habitat patches are often classified into sources and sinks based on how the subpopulations on these patches contribute to the overall metapopulation. The source-sink classification of habitat patches was first described concretely in a terrestrial context by Pulliam [11], where habitat patches were classified as sources if the local subpopulations could persist in isolation and sinks if they could not. However, this classification ignores the effect of dispersal, which is especially critical in marine metapopulations, and so Runge et al. [12] and Figueira and Crowder [13] updated the classification of source and sink patches to include both the local productivity of a patch, as well as the ability to disperse away from the habitat patch. Under this new classification a source patch is a patch on which an adult will more than self replace over the entire metapopulation and a sink is a patch on which an adult will not. Self-replacement need not occur on the same habitat patch as the adult originated, and thus under this classification a source patch may not be able to persist in isolation.

Due to the large scale larval dispersal that occurs in many marine species, it is common in marine metapopulations for sources patches to not be able to persist in isolation [14], and thus preserving a persistent metapopulation, especially in the context of MPAs, often requires more than simply preserving source patches. To maintain persistent marine metapopulations it is necessary to preserve sufficient larval exchange between closed loops of habitat patches so that an average adult can eventually self replace over multiple generations [15, 10]. If sources patches self-recruit enough larvae to persist in isolation then this can be accomplished by preserving only source patches. However if they do not it may require preserving both source and sink patches, as the sink patches may provide sufficient larval exchange back to the source patches to create a closed loop of habitat patches over which an adult can self replace. Evaluating the persistence of marine metapopulations is therefore difficult as it requires accurate measures of larval connectivity between habitat patches, as well as accurate local demographic rates of adult stages on each habitat patch [10]. Despite the difficulties, evaluating metapopulation persistence is critical in designing Marine Protected Areas in which the protected habitat patches can persist even when outside patches are exploited [16, 17, 18, 19, 20, 21].

Next-generation matrices are a useful tool that can both be used to evaluate the persistence of a metapopulation as well as identify the contribution of local habitat patches under a variety of modeling frameworks. Originally popularized in epidemiology as a simple method of calculating the basic reproduction number, *R*_0_, in compartmental models [22], next-generation matrices convert population models into generational time, so that the entries are the number of new individuals produced in each compartment, or patch in the case of metapopulations, after one generation. The individual contribution of habitat patches or evaluation of metapopulation persistence can therefore be measured for different model structures (discrete time, continuous time, etc.) under the same framework of the next-generation matrix. The column sums can be used to measure the contribution of each habitat patch over a generation and the spectral radius can be used to evaluate metapopulation persistence [23]. Next-generation operators have previously been used in ecology to calculate source and sink regions in heterogeneous environments [24, 25, 26, 23], and to evaluate the level of control required to suppress invasive species [27].

In this paper we focus on using the next-generation matrix to evaluate local patch contribution and metapopulation persistence in marine metapopulations, but the framework used here is also applicable to many other birth-jump metapopulations [28], where there is a single juvenile stage which can disperse between habitat patches and the remaining stages remain on a single habitat patch. Examples of non-marine species that exhibit this structure include plant species where seeds are caried between habitat patches [29], or insect species with a single large dispersal event such as the spruce budworm [30] or mountain pine beetle [31]. In fact, the next-generation matrix approach can even be extended to metapopulations in which adults also disperse, though the calculations become more complicated and so here we focus on species with a single dispersing stage.

Specifically, to demonstrate the utility of the method, we use the next-generation matrix to calculate the contribution of a single salmon farm to the spread of sea lice in a salmon farming region on the west coast of British Columbia. Sea lice are a parasitic marine copepod that feed on the epidermal tissues, muscles, and blood of salmon [32]. With a free living larval stage they can disperse tens of kilometres, spreading between salmon farms in a region and between wild and farmed salmon [33, 34, 35]. Lesions and stress from high sea lice infestation make adult salmon more susceptible to secondary infections, leading to large economic consequences for the salmon farming industry [36]. On wild juvenile salmon, infestation with sea lice can lead to mortality and elevated exposure to sea lice from salmon farms can contribute to population level declines in pink salmon [37]. In the context of sea lice on salmon farms we are not concerned with preserving a persistent metapopulation of sea lice parasites, but instead we use the next-generation matrix to calculate the relative contribution of each salmon farm and evaluate the effect of environmental variables on the overall growth of the sea louse metapopulation.

The specific salmon farming region that we focus on to calculate farm contribution is the Broughton Archipelago. The Broughton Archipelago is located on the west coast of Canada, between Vancouver Island and the mainland of British Columbia and has been central in evaluating the effect of sea lice from salmon farms on wild salmon [38, 35, 39, 40, 41, 37, 42, 43, 44, 45]. The area has historically had around 20 active salmon farms [46], though currently certain farms are being removed from this region in an agreement between the government of British Columbia and the Kwikwasut’inuxw Haxwa’mis, ‘Namgis, and Mamalilikulla First Nations [47]. After 2023 many of the remaining farms must be approved by both the local First Nations and the government in order to continue to operate and thus determining the farms which are acting as the largest sources of sea lice is critical during this transition period.

The paper is structured as follows. First we demonstrate how to use the next-generation matrix to calculate the contribution of local habitat patches to the metapopulation and evaluate metapopulation persistence. Next, we highlight how to construct the next-generation matrix for different types of models. Then we calculate a next-generation matrix for sea louse populations in the Broughton Archipelago to identify which salmon farms are the largest sources of sea lice in this region, evaluate the effect of the current farm removals, and investigate the effect of environmental variables on metapopulation growth. Finally, we discuss how the calculations of patch contribution and metapopulation persistence from other studies compare to the calculations using the next-generation matrix.

## 2 Materials and Methods

In this section we present details on the construction of the next-generation matrix and show how it can be used to determine the contribution of the subpopulation on a single habitat patch to the metapopulation as well as determine the persistence of the metapopulation. We then present the explicit construction of the next-generation matrix for models with age dependent demography, and present the construction for ordinary differential equation models and discrete time models in the appendix. Finally, we detail the construction of the next-generation matrix for a system of sea lice populations on salmon farms in the Broughton Archipelago.

### 2.1 Next-generation matrices

Here we construct the next generation matrix for a single species marine metapopulation with a single larval stage that can disperse between patches and where the remaining stages are confined to the habitat patch on which the larvae settle. We assume that the last stage is the only stage that produces new larvae. This assumption can be relaxed, though the entries of the next-generation matrix become slightly more complicated. We construct the next-generation matrix for models where the metapopulation is divided into *l* patches and the *m* attached stages are modelled explicitly. The larval stage is modelled implicitly, so that the birth rate into the first attached stage includes both the birth rate of larvae and the probability of larvae successfully dispersing between patches and attaching on a new patch. The lifecycle diagram for such a metapopulation is shown in Figure 1.

**Figure 1:**
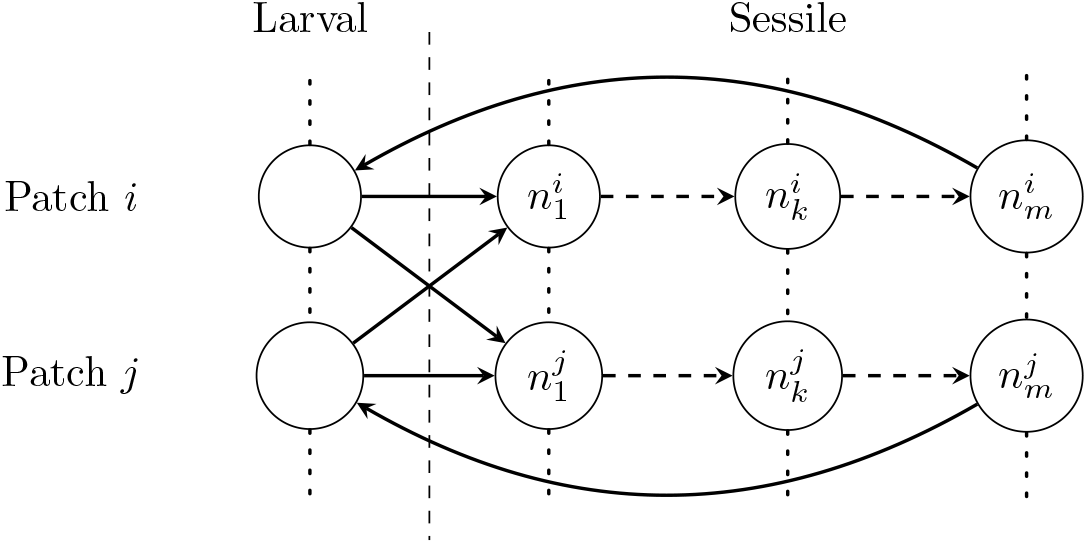
The lifecycle graph for two patches in a metapopulation of a species with a single larval stage that disperses between habitat patches and *m* sessile stages that remain on a habitat patch. The population on patch *i* in stage *k* is given by 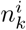.

**Figure 2:**
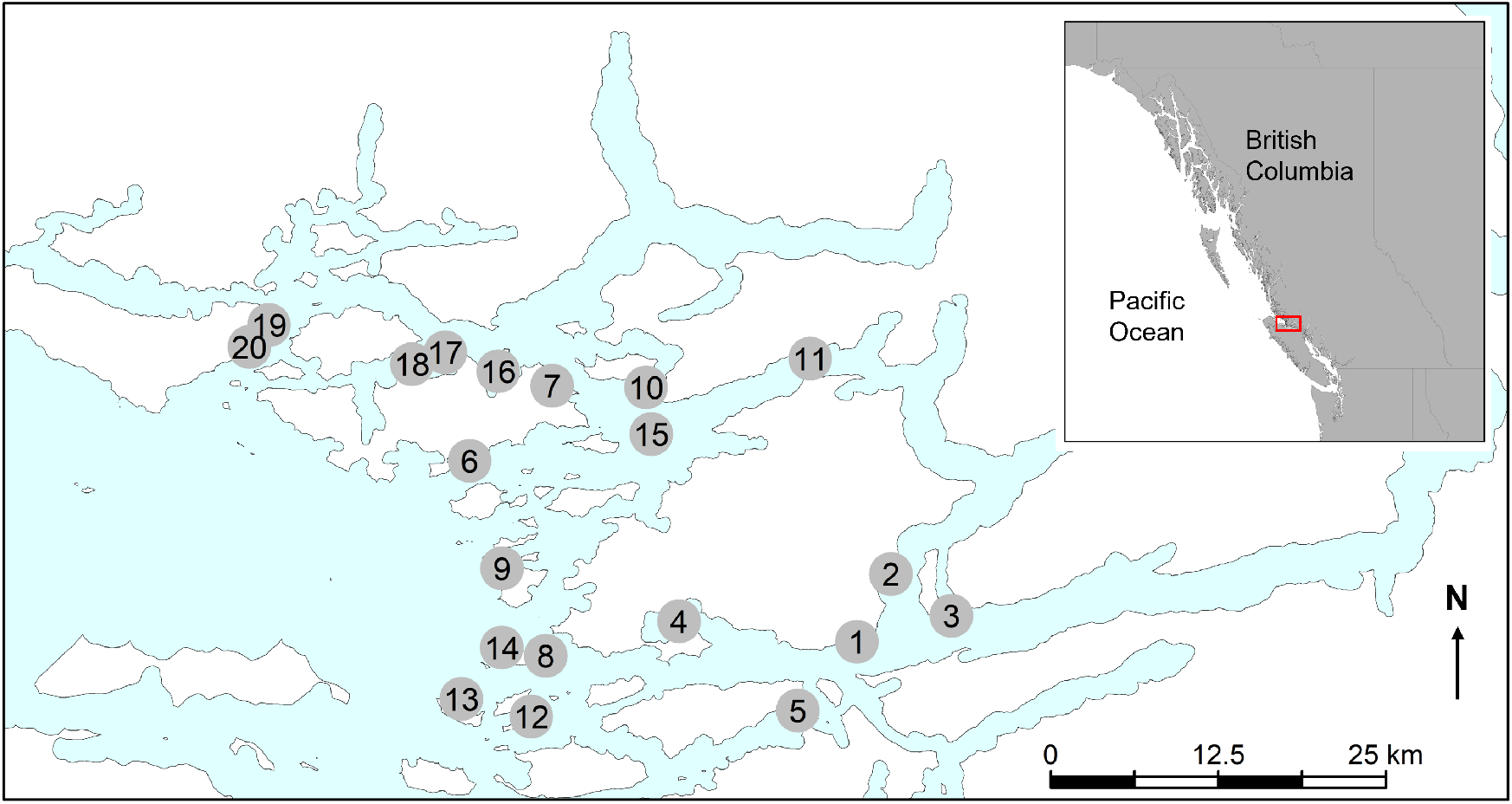
Map of the 20 historically active salmon farms in the Broughton Archipelago, for which the next-generation matrix is calculated.

We use the next-generation matrix to calculate local patch contribution to the metapopulation in the context of low population density. To calculate the next-generation matrix it is necessary to linearize a potentially density dependent model around the zero equilibrium so that the effect of density dependence at higher population sizes is ignored when calculating patch contribution. This approach of ignoring density dependence is common when determining persistence or patch contribution of marine metapopulations, as the focus is either on determining if a metapopulation can persist at all, or determining which habitat patches are acting as population sinks and which are acting as population sources [15, 10, 24, 23]. Alternatively it is also useful for determining patch contribution in metapopulations of species that are being actively controlled to remain at low densities, such as sea lice on salmon farms, which is our focus in section 2.4. Another common assumption in the theory of persistence also made here is that there is no discernible Allee effect in any of the patches [15, 24, 10].

Under any model structure, the element in row *i* and column *j* of the next-generation matrix gives the number of new individuals produced on patch *i* by one new initial individual on patch *j*. However, how ‘new’ individuals are defined is subject to interpretation. Here, we construct the next-generation matrix under modeling frameworks that only consider the sessile stages explicitly, and so ‘new’ individuals will be newly attached stage 1 individuals. Then, when the number of new individuals on patch *i* produced from one new individual on patch *j* are tracked, the new individual must first survive and reproduce on patch *j*, before larvae disperse and arrive on patch *i*. In this way patch contribution, as will be calculated in section 2.2, is primarily a function of the local patch demography which is then coupled with dispersal to other patches. Under this framework, the entries of the next-generation matrix, *K*, for all model structures can be given by:

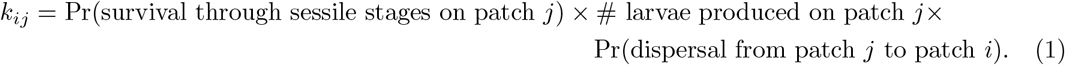

However if the next-generation matrix is constructed for models that explicitly model the larval stage, then the larval stage is often considered the first stage. The next-generation matrix will be slightly different in this case as well as the calculation of patch contribution, though the calculation of metapopulation persistence will be the same. We illustrate the differences between constructions of the next-generation matrix in the Discussion and show how the calculation of persistence remains the same under all constructions.

### 2.2 Determining patch contribution and metapopulation persistence

Here we show how to use the next-generation matrix, *K*, to determine the contribution of each patch to the metapopulation and evaluate metapopulation persistence. To determine the contribution of a specific patch to the metapopulation we track the total number of new individuals produced across the metapopulation after one generation from an initial individual starting on that patch. The entries of the next-generation matrix, *k*_*ij*_ give the number of new individuals produced on patch *i* from an initial individual on patch *j*. Therefore if we define

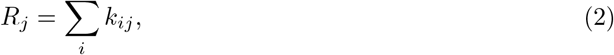

so that *R*_*j*_ is the *j*th column sum of *K*, then *R*_*j*_ is the total number of new individuals produced across all patches from an initial individual starting on patch *j* and can be used to define the contribution of patch *j* to the entire metapopulation.

This definition of patch contribution easily lends itself to classifying local habitat patches as population sources or sinks. If *R*_*j*_ *<* 1, then an individual on patch *j* cannot replace itself over the entire metapopulation, and thus patch *j* is defined as a sink [23]. If *R*_*j*_ *>* 1 then one individual on patch *j* is producing more than one individual over the entire population and so patch *j* is defined as a source. To calculate persistence we can use the basic reproduction number, *R*_0_, which can be calculated as

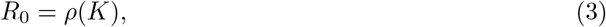

where *ρ*() is the spectral radius. If *R*_0_ *>* 1 then the metapopulation will persist and if *R*_0_ *<* 1 then the metapopulation will go extinct, a relationship which holds under any of the model formulations considered here [22, 23, 48, 49]. The only condition required is that *K* be irreducible, which is biologically satisfied if there is some small positive probability that larvae leaving one patch can eventually arrive on any other patch.

There are several biologically reasonable properties that also exist mathematically under this framework. First, if the population on any single habitat patch can persist on its own, so that *k*_*ii*_ *>* 1, then the entire metapopulation will persist and *R*_0_ *>* 1 [23]. Second, a metapopulation consisting only of sink patches cannot persist and a metapopulation consisting only of sources cannot go extinct. The mathematical underpinning of these relationships is that the spectral radius must be between the minimum and maximum column sums of a matrix, so in terms of our metapopulation quantities

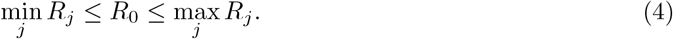

Having defined patch contribution and persistence in terms of the next-generation matrix, we now demonstrate how to calculate the next-generation matrix for a model with age dependent demography, as this is the modeling structure commonly used for sea lice on salmon farms. We also present the construction of the next-generation matrix for discrete time models and ordinary differential equation models in the Appendices A and B, so that the details are present for the most commonly used population models

### 2.3 Calculating the next-generation matrix for models with age dependent demography

Here we calculate the next-generation matrix for models which allow for the maturation, survival, birth, and dispersal rates to depend not only on the stage and patch location of an individual, but also on the time that they have spent in a stage. This time is often referred to as stage-age and so in these models there are two time variables: the global time of the system, *t*, and the time that an individual has spent in a particular stage, their stage-age, *a*. These models can be specified either as McKendrick von-Foerster partial differential equations, integrodifferential equations, or renewal equations [50]. In all cases, the dependence of the maturation rate on time spent in a stage allows for the addition of more realistic maturation functions where most individuals mature at some intermediate stage-age, or after some minimum time spent in the stage. In contrast, if models are formulated using ordinary differential equations the time in a stage is always exponentially distributed.

However, the specification of the model equations for these models can be rather complicated and so here we construct the next-generation matrix directly from the maturation, survival, birth, and dispersal functions, but the full model and the derivation of the next-generation matrix can be found in Appendix C as well as in [23]. Essentially the construction involves tracking the probability that an individual survives through the different stages on a specific patch, the number of larvae that they produce, and the probability that the larvae successfully disperse from one patch from another. If the probability of survival in stage *k* on patch *j* at stage-age *a* is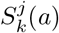, the maturation rate from stage *k* to *k* + 1 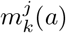, the birth rate of larvae on patch *j* is *b*^*j*^(*a*), and the probability of a larva leaving patch *j* and successfully attaching on patch *i* is *p*^*ij*^, then the entries of the next-generation matrix, *K*, are given by

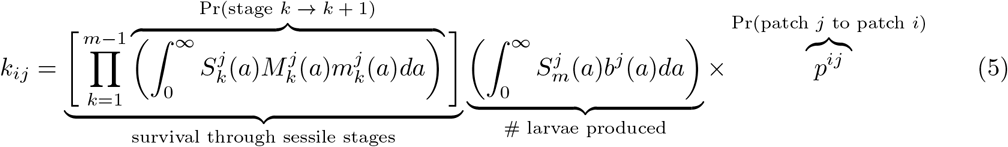

where 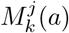 is the probability that an individual has not yet matured from stage *k* to *k* + 1 and can be calculated as 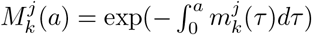.

### 2.4 Application: sea lice on salmon farms in the Broughton Archipelago

In this section we construct a next-generation matrix to determine the contribution of a single salmon farm to the spread of sea lice in a salmon farming region on the west coast of British Columbia, the Broughton Archipelago. To determine the level of sea lice dispersal away from salmon farms, we use a hydrodynamic model, coupled with a particle tracking model, to track sea lice particles released from 20 historical farms in the Broughton Archipelago [51]. The coupled hydrodynamic and particle tracking models have been validated with sea louse counts on salmon farms in the Broughton [52]. We construct one next-generation matrix for the 20 historical farms in the region for which the hydrodynamic particle tracking model was run, as well as one for the 11 remaining farms in the area after 2023, subject to First Nations and governmental approval [47].

#### 2.4.1 Modeling framework

Sea lice maturation through stages is often modelled with stage-age dependent maturation functions [53, 54, 55] and thus we construct the next-generation matrix for sea lice using age dependent demography, as shown in section 2.3.

#### 2.4.2 Dispersal

The probability that a sea louse larvae leaves from one farm and successfully arrives on another depends on several factors including ocean current, temperature, and salinity. To accurately capture this probability it is necessary to use a computational hydrodynamic model that can track the spread of larvae originating from a given farm as well as the dependence of larval survival on temperature and salinity. In order to determine the probability of larvae dispersing between farms we use connectivity matrices from Cantrell et al. [51]. Details on these connectivity matrices can be found in [51]. Briefly, these connectivity matrices are calculated by applying Kernel Density Estimation (KDE) to particle tracking simulations to calculate the infectious density of sea lice at each farm, originating from a given farm. The particle tracking simulations are run on output generated by a Finite Volume Community Ocean Model (FVCOM) which uses data on tides, wind surface-heating, and river discharge to simulate three-dimensional ocean velocity, temperature, and salinity [56]. In the particle tracking simulation the survival of sea louse particles is dependent on temperature and their maturation from non-infectious to infectious larvae is dependent on temperature.

The infectious densities of sea lice are calculated for each particle release day by taking daily snapshots of the particle locations of infectious lice for 11 days post release and then applying Kernel Density Estimation to the accumulated daily snapshots. A connectivity matrix is then calculated for each particle release day, where the entry in row *i* and column *j* of the connectivity matrix is the infectious density of larvae over farm *i*, produced by larvae initially leaving farm *j*. In [51] the infectious densities were calculated from 24 hours of particle releases, where 50 particles were released each hour and so to calculate the infectious density of one initial release particle we divide the entries in each connectivity matrix by 1200 (50 × 24). Then, to create a single connectivity matrix, *C*, for the 20 farms in the Broughton Archipelago we take the average over all the connectivity matrices created for particles released between March 14th and July 20th, 2009.

The necessary quantity to construct the next-generation matrix is *p*^*ij*^, the probability that larvae leaving farm *j* will successfully attach on farm *i*. To estimate *p*^*ij*^ from the entries of the connectivity matrix, *c*_*ij*_, there are several assumptions that need to be made. If we assume that the number of lice that arrive onto farms is small compared to the total number of lice in the water column, so that lice arriving onto farms do not significantly affect the density of lice in the water column, then

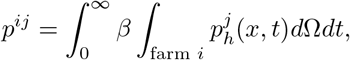

Where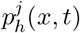 is the two dimensional density of infectious lice produced from farm *j* that are still alive in the water column at position *x* and time *t*, and *β* is the arrival rate of lice moving over the farm arriving onto the farm. However the entries of the connectivity matrix, *c*_*ij*_, are the infectious density of larvae over farm *i*, produced by larvae leaving farm *j*. The infectious densities are calculated by applying Kernel Density Estimation to daily snapshots of infectious particles over 11 days, starting at time *t* = 0, so roughly

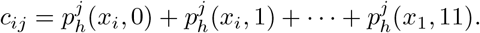

We can therefore roughly calculate *p*^*ij*^ from *c*_*ij*_ by assuming that the integrals over time and space can be approximated using their Riemann sums, and that the area of a farm is roughly 0.01 km^2^, where

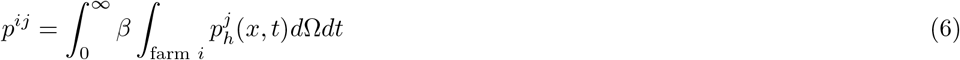

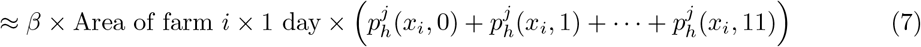

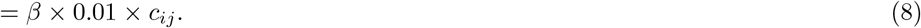

Therefore to calculate *p*^*ij*^ from *c*_*ij*_ we need to estimate the arrival rate *β*. However, very little is known about the arrival rate of lice dispersing from one farm to another, and so the estimate presented here is very uncertain and could be orders of magnitude off from the true arrival rate. We assume here that *β* = 100*/*day, and thus the average waiting time for infectious sea lice in the water column surrounding the farm to arrive on the farm is roughly 15 minutes (1*/β* = (1 day*/*100) × (24 hrs*/*1 day) × (60 minutes*/*1 hr) = 14.4 minutes). However the waiting time could be as little as 1 minute or as long as 1 hour, depending on environmental conditions. Moreover, this is ignoring the fact that some infectious lice may never attach, and the estimates of the proportion of lice which successfully attach at all varies from 80% to 0.5% under different lab conditions [57].

It can be seen from equations (5) and (8) that each entry in the next-generation matrix, *K*, is a linear function of the arrival rate *β*, and therefore changing *β* will not affect the relative ordering of the contributions of different salmon farms, given by *R*_*j*_ (equation 2), it will only affect the absolute magnitude. The basic reproduction rate, *R*_0_ = *ρ*(*K*), is also a linear function of *β* and therefore any increase or decrease in *β* will result in the same proportional increase or decrease in *R*_0_. However, the purpose of this application is not to accurately estimate the basic reproduction number, *R*_0_, for the Broughton Archipelago, or to accurately estimate the contribution of an individual farm, *R*_*j*_, but rather to compare the relative contributions of different salmon farms in the system, and to investigate the effect of environmental variables on the basic reproduction number. Therefore we present our estimate of the arrival rate for these purposes only, and our value of *R*_0_ found in the results should not be taken as an accurate estimate.

#### 2.4.3 Demography

Once infectious sea lice larvae attach to their salmonid hosts they must survive and mature through several attached life stages before they can produce offspring. These demographic rates are dependent on salinity and temperature, and thus some salmon farms may be more productive than others due to favourable environmental conditions. To capture the dependence of demography on salinity and temperature we simplify the attched sea lice life cycle down to three main stages: chalimus, pre-adult, and adult. Survival in each stage is salinity dependent, maturation is temperature dependent, and egg viability and production depends on both salinity and temperature. The demographic functions that we use are from models which have previously been fit to sea louse population data and are shown in Table 1.

**Table 1:**
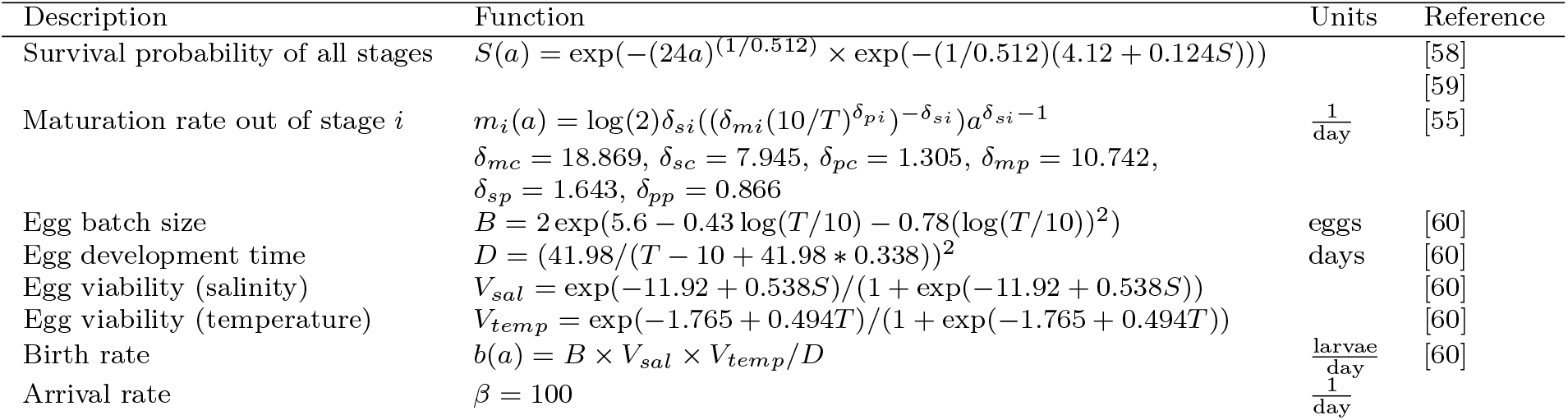
The maturation, survival, and birth functions used to create the next-generation matrix for sea louse populations on salmon farms in the Broughton Archipelago. The sea louse life cycle is simplified to three attached stages: chalimus, pre-adult and adult. For the maturation rate out of stage *i, i* ∈ (*c, p*), where *c* refers to the chalimus stage and *p* refers to the pre-adult stage.

We calculate the on-patch component of the elements of next-generation matrix, *k*_*ij*_, by integrating the demographic functions over all time,

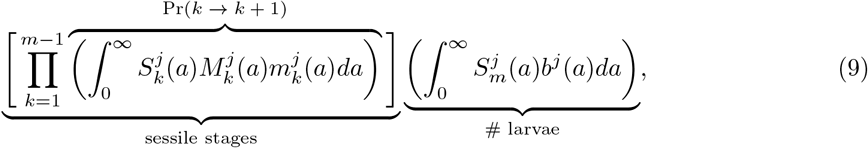

and refer to this as the productivity of patch *j*. To determine the specific temperature and salinity dependent demographic rates at each farm we find the temperature and salinity that each sea louse particle experiences in the particle tracking simulation when initially released from a farm. We then use the average temperature and salinity of particles over all releases.

We also investigate the effect of varying temperature and salinity on the relative growth and persistence of the metapopulation. To do so, we must calculate survival and maturation rates for temperatures and salinities that farms may not experience in the period for which the FVCOM was run. To keep the variability of temperatures and salinities that exists between farms, we multiply the new temperature or salinity at which we want to evaluate persistence, by the ratio of the mean farm temperature or salinity divided by the mean total temperature or salinity experienced by all farms.

## 3 Results

In this section we present the next-generation matrix for sea lice populations on salmon farms in the Broughton Archipelago, the construction of which is detailed in section 2.4, and determine the relative patch contribution of each farm. We use this system to highlight to potential differences between the connectivity matrix, which only contains information surrounding the probability of dispersal from one farm to another, and the next-generation matrix, which combines dispersal between farms and local productivity of sea lice on a salmon farm. We then demonstrate how the next-generation matrix can be used to investigate the effect of changing demographic rates on growth and persistence in this system. Finally, in the context of salmon farm removal from the Broughton Archipelago, we investigate how the removal of habitat patches affects patch contribution and persistence in this sea louse metapopulation.

The next-generation matrix for sea lice populations on salmon farms in the Broughton Archipelago is shown in Figure 3a). The patch contribution of each salmon farm is given by *R*_*j*_, the *j*th column sum, which is presented at the bottom of each column. We also present the row sums to the left each row, to identify which farms are receiving the most sea lice from other farms in the region. Farm 2 is the largest contributor of sea lice to the metapopulation, followed by farms 17 and 18, whereas farm 12 has the lowest contribution. The farms receiving the most sea lice, in declining order, are farms 3, 16, and 7.

**Figure 3:**
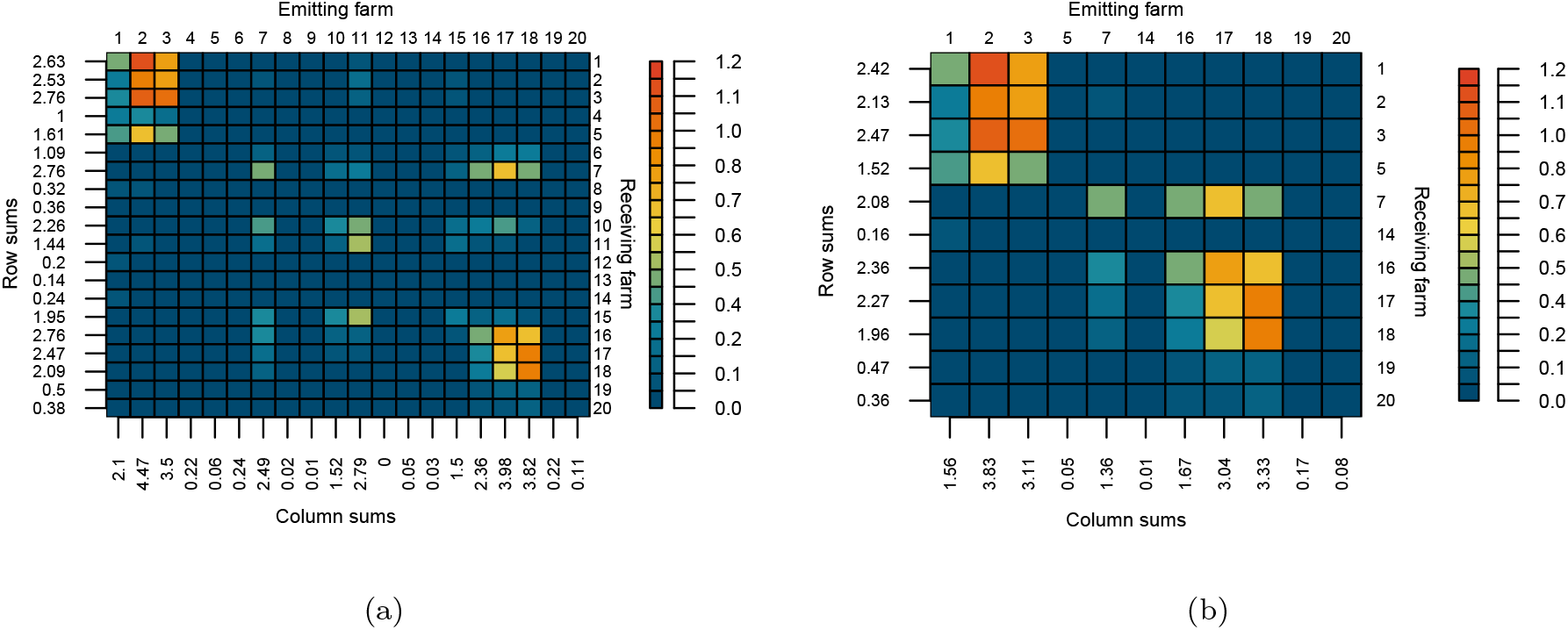
a) The next-generation matrix for the 20 historically active farms in the Broughton Archipelago, and b) the next-generation matrix containing only the farms remaining in the Broughton Archipelago after 2023, subject to First Nations and governmental approval [47]. The entries of the next-generations matrices, *k*_*ij*_ are the number of new chalimus stage lice produced on farm *i* from one initial chalimus on farm *j*. The column sums, *R*_*j*_, are the total number of chalimus produced on all farms from an initial chalimus on farm *j* and are shown below each column. Likewise the row sums are the number of new chalimus received by each farm from all other farms and are shown on the left of each row. These numbers should be taken as relative, rather than absolute, as we do not have a very accurate estimate for the arrival rate of sea lice onto farms, *β*.

To better understand the details and construction of the next-generation matrix, we also present the connectivity matrix for this system in Figure 4 and the productivity (total number of new larvae produced from one attached chalimus louse) of each farm in Table 2. The (*i, j*)th entry of the connectivity matrix, *c*_*ij*_, is the infectious density over farm *i* of lice leaving farm *j* and the (*i, j*)th entry of the next-generation matrix is constructed by multiplying the productivity of farm *j* (equation 2.4.2) *p*^*ij*^ = *β* × 0.01 × *c*_*ij*_, the probability that a larvae leaving farm *j* attaches on farm *i* (8). The farms with the largest column sums of the connectivity matrix are, in declining order, farms 18, 2, and 17. However, farm 2 has a higher productivity than farms 17 or 18, and when the productivity of farm 2 is multiplied by the connectivity then it becomes the largest source of sea lice in the region, as identified by the next-generation matrix.

**Table 2:** The number of new larvae produced on each farm by a single louse starting in the chalimus stage. The first row is the farm number and the second row is the number of larvae produced.

|  |  |  |  |  |  |  |  |  |  |  |
| --- | --- | --- | --- | --- | --- | --- | --- | --- | --- | --- |
| Farm number | 1 | 2 | 3 | 4 | 5 | 6 | 7 | 8 | 9 | 10 |
| Number of larvae produced | 1355 | 1582 | 1462 | 239 | 244 | 612 | 1064 | 111 | 56 | 1060 |
| Farm number | 11 | 12 | 13 | 14 | 15 | 16 | 17 | 18 | 19 | 20 |
| Number of larvae produced | 1529 | 33 | 331 | 175 | 1255 | 1143 | 1483 | 1348 | 584 | 464 |

**Figure 4:**
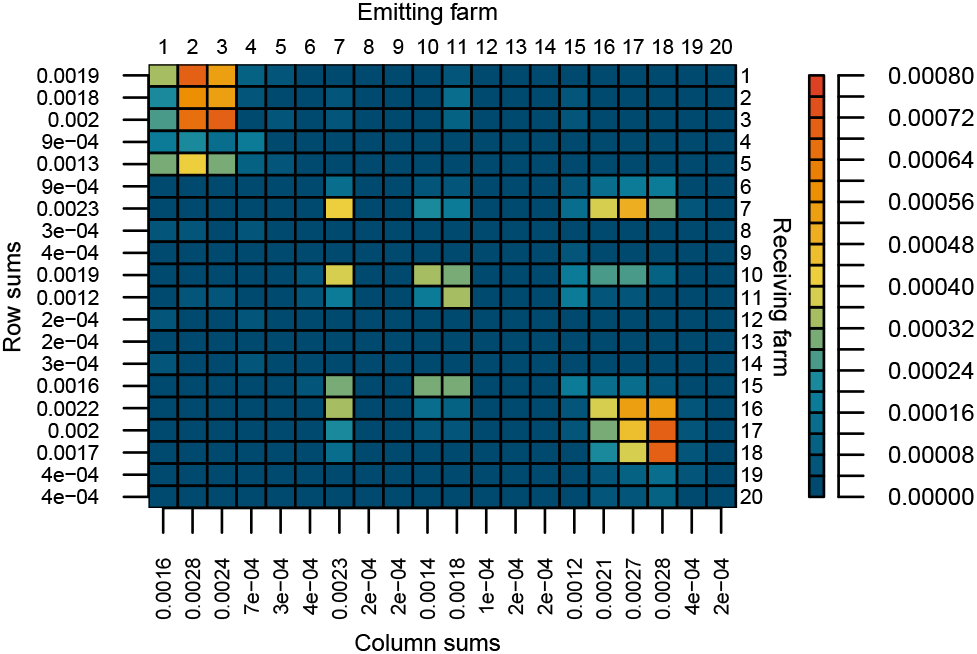
The connectivity matrix, *C*, for sea lice larvae dispersing between salmon farms, created by averaging all connectivity matrices from [51] over the simulation period. The (*i, j*)th entry is the infectious density of larvae (1*/*km^2^) over farm *i* that have left from farm *j*. Column and row sums are shown below and to the left of each column and row, respectively.

A further look into the connectivity matrix and productivity table provides more insight into the underlying drivers of the contribution of each farm to the sea louse population. Many of the farms with low connectivity also have low productivity, which may be due to the fact that temperature and salinity affect on farm demographic rates as well as survival and maturation in the particle tracking simulation which underlies the connectivity matrix. However, there are certain farms, such as 11 and 15, which have a comparably high productivity compared to their connectivity. These farms are located in favourable environments with respect to temperature and salinity, but low connectivity due to either distance from other farms or unfavourable currents prevents these farms from acting as larger sources of sea lice.

Here we also examine how temperature and salinity affect the overall growth and relative persistence of the sea louse metapopulation, as shown in Figure 5. The persistence of the metapopulation is determined by the basic reproduction number, *R*_0_, which is calculated as the spectral radius of the next-generation matrix. As we do not have an accurate estimate for *β*, we examine the effect of temperature and salinity on the relative change in persistence or growth of the metapopulation, but refrain from commenting on the absolute growth, as measured by *R*_0_. We can see that as both temperature and salinity increases, the overall growth of the metapopulation increases, and that salinity has a larger effect on metapopulation growth than temperature. What is also interesting, but cannot be seen from the figure, is that as salinity increases, the farm that receives the most lice switches from farm 3 to farm 7.

**Figure 5:**
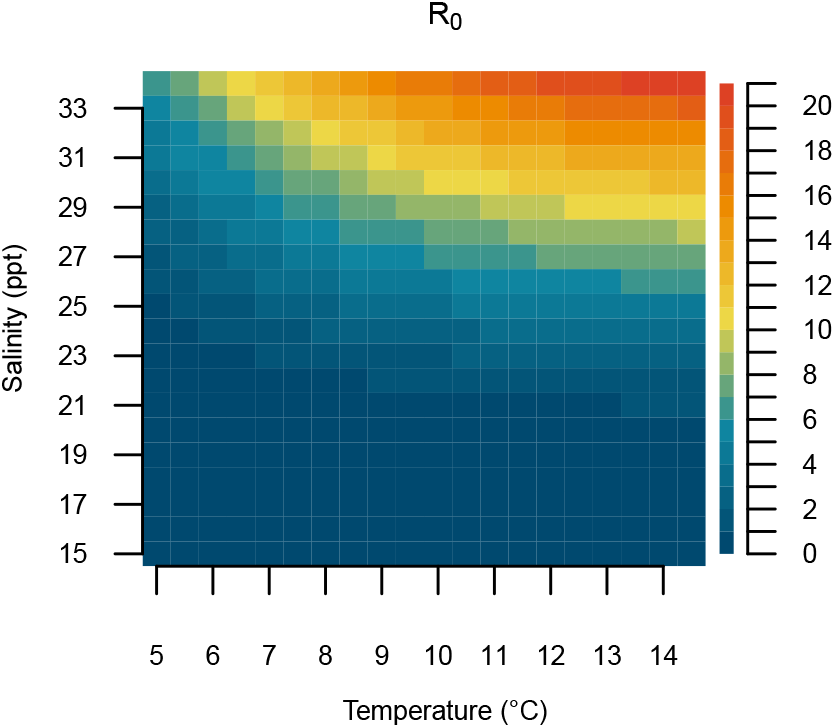
The effect of temperature and salinity on the overall growth or persistence of the original sea lice metapopulation of 20 farms, as described by the basic reproduction number, *R*_0_. We do not have a good estimate for the arrival rate of sea lice onto farms, *β*, and so the *R*_0_ values should only be interpreted relative to each other, rather than as absolute values.

In light of the removal of salmon farms in the Broughton Archipelago we also create a nextgeneration matrix consisting only of the farms which will remain after 2023 subject to First Nations and government approval, shown in Figure 3b), and examine the differences between this matrix and the next-generation matrix with all farms. Most of the farms which are acting as the largest sources of sea lice in this region remain, with the exception of farm 11, the fifth largest source in the original network, which has now been removed. Farm 11, while not the largest source, did have the largest betweenness score based solely on the connectivity matrix [51], and thus may have been acting as a connecting farm between the two large clusters of source farms. However, since none of the other large source farms have been removed, the overall growth of the metapopulation has only decreased from *R*_0_ = 2.33 (original next-generation matrix) to *R*_0_ = 2.25. Again these numbers are calculated using a very rough estimate of the arrival rate onto farms, detailed in section 2.4.2, and thus it is their relative similarity that is important, rather than the absolute magnitude.

## 4 Discussion

In this paper we demonstrated how to use the next-generation matrix to calculate the contribution of each habitat patch to the metapopulation and measure the overall persistence of a metapopulation. We detailed the construction of the next-generation matrix under different model structures to demonstrate the breadth of the approach to several systems. We then constructed the next-generation matrix under an age dependent modeling framework for sea lice populations on salmon farms in the Broughton Archipelago to illustrate how this approach can be applied to a real system. We determined which salmon farms may be acting as the largest sources of sea lice in this region, how the metapopulation will change once certain farms are removed, and examined the effect of temperature and salinity on the relative growth and persistence of this metapopulation.

Next-generation matrices have been used extensively in epidemiology to study the spread of infectious diseases but have recently been introduced in ecology [24, 61, 26, 27] and evolutionary analysis [62]. One of the key benefits of using next-generation matrices in epidemiology is that the basic reproduction number, *R*_0_, for a disease can be calculated as the spectral radius of the nextgeneration matrix, which is often much simpler than calculating the eigenvalues of the full system to determine spread. In ecology, one main advantage of this approach is that the mathematical calculation of *R*_0_ can be broken down into biologically relevant quantities, for example the contribution of different dispersal pathways to growth in a population [63] or the contribution of populations on different habitat patches [23]. While not novel, we hope next-generation matrices can be used more frequently as a simple and easily biologically interpretable method to measure the contribution of local habitat patches to a metapopulation and determine overall persistence.

We are also by no means the first to attempt to calculate the contribution of a local population, classify patches into sources and sinks, or attempt to measure the persistence of metapopulations. In the context of low densities Pulliam [11] defined a source as a habitat patch that would grow in the absence of immigration and emigration and a sink as a habitat patch that would decline in the absence of immigration and emigration. This is similar to only using the entries along the diagonal of the next-generation matrix to classify sources or sinks, except growth or decline was measured after one time step in a discrete time model, rather than one generation. However, as discussed at the end of section 2.2, it is possible to have a metapopulation composed only of sinks based on this definition (*k*_*ii*_ *<* 1 for all *i*), that persists.

Recognizing that dispersal between patches should also considered when classifying habitat patches as sources or sinks, both Runge [12] and Figueira and Crowder [13] defined new metrics to classify habitat patches that track the contribution of adults on a patch in one time step to the total population on all patches in the next timestep. These metrics are similar to our patch contribution metric *R*_*j*_, except they measure the contribution over one time step rather than one generation, similar to using the dominant eigenvalue of the projection matrix *A* to determine the stability of the discrete system *n*(*t* + 1) = *An*(*t*), rather than the spectral radius of the next-generation matrix *K*. However, the calculations can become complicated if the population is stage structured and *A* is large (Appendix A, [12]) and the metrics do not easily generalize to systems described by ordinary differential equations.

There are other measures of persistence in metapopulations which do track the number of new individuals contributed to the metapopulation after one generation from an initial individual on one patch, though they use different starting stages for the initial and new individuals. Krkošek and Lewis [24] define a next-generation operator for general heterogeneous populations which tracks the number of new adults produced in the population from one initial adult after one generation. If *b*_*j*_ is the reproductive output on patch *j, p*_*ij*_ is the probability of larvae dispersing between patch *j* successfully arrives on patch *i*, and *a*_*i*_ is the survival to adulthood on patch *i*, then the contribution of patch *j* to patch *i* according to Krkošek and Lewis [24] can be calculated as *b*_*j*_*p*_*ij*_*a*_*i*_. This patch contribution metric cannot be calculated directly from the next-generation matrices shown in this paper unless there is only one stage. Burgess et al. [10], following Hastings and Botsford [15], track the number of new larvae on all patches produced by an initial larvae. The entries of their ‘connectivity matrix’ (similar to our next-generation matrix), *c*_*ij*_, are given by *p*_*ij*_*a*_*i*_*b*_*i*_. The entries of their connectivity matrix would be the entries of the next-generation matrix if it was calculated from a model with an explicit larval stage. However under this construction the contribution of patch *j*, as calculated by the *j*th column sum of their connectivity matrix, is primarily a function of the demography of the patches on which larvae dispersing from *j* settle, rather than the local patch demography of patch *j* itself, as it is when using the next-generation matrix as we have formulated it in this chapter.

The contribution of each habitat patch to the metapopulation will depend on the stage at which the generational output is measured, but the persistence of the metapopulation is equivalent under all of these frameworks. This is because there is only one component of the life cycle where movement can occur between patches (larval stage) and at all other stages individuals remain on a patch. Let *P* be a matrix with entries *p*_*ij*_, *B* be a diagonal matrix with entries *b*_*j*_ and *A* be a diagonal matrix with entries *a*_*j*_, where *p*_*ij*_, *b*_*j*_ and *a*_*j*_ are the same as the preceding paragraph. If we measure generational output starting at the first attached stage, as we do in this chapter, the next-generation matrix can be written as *PAB*, if we measure generational output starting at the larval stage according to Burgess et al. [10], the matrix can be written as *ABP*, and if we measure generational output starting at the adult stage according to Krkošek and Lewis [24] then the matrix can be written as *APB*. This is because when *P* is multiplied by a diagonal matrix on the right, the entries of the diagonal matrix multiply each column of *P* and when *P* is multiplied on the left, the entries multiply each row. Now for any two matrices *X* and *Y, XY* and *Y X* have the same eigenvalues, and because matrix multiplication is associative each of the matrices *PAB, ABP*, and *APB* all have the same eigenvalues as well, and therefore also the same spectral radius. Therefore in any of the formulations the metapopulation will only persist if *ρ*(*PAB*) = *ρ*(*ABP*) = *ρ*(*APB*) *>* 1.

While the metapopulation persistence criteria is equivalent to other formulations [24, 10, 15] we believe using the next-generation matrix provides several advantages. First, the framework is the same for discrete time, continuous time, and age structured systems of equations. Second, it is easy to convert between the full system of equations and the next-generation matrix, thus if a system of equations has already been parameterized for a given region, it is easy to calculate the contribution of each habitat patch using the next-generation matrix. Third, there is natural extension of the next-generation matrix to systems where multiple stages can reproduce, or when adults can migrate between patches. In discrete time and continuous time the composition of the matrix in terms of the fecundity and transition matrix still holds, only the formulas given in terms of the survival, maturation, and birth rates no longer hold. The next-generation matrix can also be extended as an operator to integro-difference equations [24, 26] if space is continuous, rather than divided into discrete habitat patches. Lastly, in continuous time systems it is possible to connect the patch contribution computed from the next-generation matrix, *R*_*j*_, and the transient dynamics of the metapopulation (Harrington et al, in press).

In our application of the next-generation matrix to sea lice populations on salmon farms in the Broughton Archipelago, we believe there are two main sources of uncertainty when calculating the entries of the next-generation matrix: uncertainty in the arrival rate, *β*, and uncertainty in the number of larval sea lice that remain inside the salmon farm in the particle tracking model. In lab studies, the proportion of infective copepodid lice that attach to salmon varies enormously, from as low as 0.5% to as high as *>* 80%, with recent estimates between 12 − 56% depending on temperature [57]. Moreover, the time during which copepodids are given to attach in these experiments also varies, from a few seconds, to 8 hours. If we assume an exponential waiting time model, *P* = 1 −*e*^*−βt*^, where *P* is the proportion of copepodids that have attached to salmon, then these lab studies give estimates of *β* between 3/day and 47/day, which are lower than the estimate of 100/day which we use in this paper. Any estimates less than 44/day would result in a basic reproduction number, *R*_0_ *<* 1, which would mean that the overall sea louse population is not increasing.

However, there is abundant evidence that in the Broughton Archipelago sea lice populations on many salmon farms will grow exponentially in the absence of treatment by farm managers [64, 65, 66, 52, 67]. This discrepancy may be due to the fact that while the hydrodynamic and particle tracking models give good estimates of sea louse dispersal between farms [52], they likely underestimate the number of sea louse larvae that remain inside a salmon farm. In the hydrodynamic model the location of the farm does not alter the local hydrodynamics that occur within the farm, but in reality the the presence of salmon swimming within the net pens may alter the local hydrodynamics sufficiently that some proportion of larvae remain within the farm. To correct for this, it would be possible to add a diagonal matrix to the next-generation matrix presented in this paper, where each entry on the diagonal is the productivity of farm *j*, multiplied by the number of larvae which remain within and successfully attach to salmon on the farm. Because the most productive farms are also the most highly connected in the Broughton, this would likely not change the relative contribution of each farm, but would increase the absolute contribution of each farm as well as *R*_0_.

In addition to the sources of uncertainty in the next-generation matrix, there are some other technical aspects which should be considered when applying the next-generation matrix to calculate patch contribution of metapopulation persistence in specific systems. In order to use the next-generation matrix to calculate patch contribution or persistence we are assuming that our system is autonomous, and that the demographic rates do not change with time. In reality for most systems, including sea lice on salmon farms, environmental variables will fluctuate over time, potentially changing the demographic rates of the population [68]. In our case temperature and salinity change over the course of the spring, but we calculate the next-generation matrix using the mean temperature and salinity that sea lice experience during the particle tracking simulation window. Therefore the entries in our next-generation matrix may be slightly different than the true number of newly attached lice produced on other farms from one initially attached louse, depending on the exact time that the louse began its lifecyle during the spring. For temporally oscillating systems it is possible to correct for this difference, though the entries of the next-generation matrix no longer have a simple form and must be computed computationally [59].

Over time periods where temperature and salinity are relatively constant, we can use the next-generation matrix to examine the overall effect of environmental variables on the growth of the metapopulation. The demographic rates at each stage depend explicitly on temperature and salinity but alone, or in a full system of equations, it can be difficult to examine the overall effect of changing environmental variables. However, the basic reproduction number *R*_0_, calculated from the spectral radius of the next-generation matrix, provides a useful metric of the overall effect. We can infer how the growth of the metapopulation may change among seasons, or as the ocean warms. For sea lice on salmon farms in the Broughton Archipelago the effect that of temperature and salinity on *R*_0_ is very similar to previous results found for a single farm [59]. With updated temperature and salinity at each farm, we could calculate the change in growth over years in the springtime, which may help explain the recent sea louse outbreaks during warm years in the Broughton Archipelago [66]. When examining the effect of temperature and salinity on *R*_0_ we do not rerun the hydrodynamic and particle tracking models to recreate the connectivity matrices under new temperature and salinity scenarios, as this is very computationally intensive, but we expect connectivity to increase as temperature and salinity increases due to higher survival of sea lice and faster maturation. However, we believe it would be valuable to rerun hydrodynamic models under different projected ocean scenarios, to investigate the precise changes in connectivity that may occur.

Specific to management of sea lice populations on salmon farms in the Broughton Archipelago, there are several insights to be gained from our results. The first is that the farms that in the most productive environments are also the most highly connected, and thus become the largest contributors of sea lice to this sea louse metapopulation. They occur in two main clusters (shown in Figure 3) and both clusters of farms will remain in the Broughton Archipelago in the current removal plan, subject to First Nations and governmental approval [47]. It should be noted that these farms may not necessarily be producing the most number of lice compared to other farms in the region at a given time if their louse population is currently lower than other farms, rather they are the farms that have the largest potential to contribute to spread when sea louse numbers are even across farms. However, due to the highly connected nature of these clusters, coordinated treatment between farms in the clusters or all farms in the region could reduce the number of treatments required and number of sea lice produced on all farms [69]. An interesting avenue of future research would be to connect the productivity of the remaining farms in the Broughton Archipelago with the Kernel Density Estimates of sea louse dispersal from Cantrell et al. [51] to measure the exposure of migrating wild salmon to sea louse infection from these farms.

## 5 Acknowledgements

The authors would like to thank the members of the Lewis lab for many helpful discussions and suggestions. PDH gratefully acknowledges an NSERC-CGSM scholarship, Queen Elizabeth II scholarship, and an Alberta Graduate Excellence Scholarship; and MAL gratefully acknowledges the Canada Research Chair program and an NSERC Discovery Grant.

## A Calculating the next-generation matrix for differential equation models

Here we briefly formalize the calculation of the next-generation matrix for differential equation models. Again, we are considering a model for a species with *m* sessile stages on *l* patches with a larval stage that can disperse between patches and all other stages are sedentary and confined a patch. Let 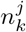 be the population in sessile stage *k* on patch *j*, then the population dynamics at low population densities can be described by

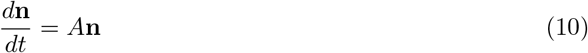

where **n** is a population vector of size *m* ×*l* describing the population size on all patches at all stages and *A* is a matrix describing the population dynamics at low population densities. We model the population size at low densities because we are interested in identifying which patches are acting as sources and supporting the population at low densities and which are acting as sinks. Therefore equation 10 should be thought of as the linearization about the zero equilibrium of a potentially more complex model that may include density dependence. The elements of *A* contain the maturation and death rates of sedentary stages confined to patches as well as the rates of larval dispersal and attachment between patches.

The next-generation matrix for this system can be calculated by first decomposing *A* = *F* − *V* where *F* is a non-negative matrix with entries that describe the rates of larval birth and probability of dispersal between patches and attachment as the first sedentary phase, and *V* is a non-singular M matrix [70] with entries that describe the maturation rates between stages and death rates in a stage [22]. Because *V* is a non-singular M matrix, *V* ^*−*1^ is non-negative. Following van den Driessche and Watmough [22] with notation from Diekmann et al. [71], the next-generation matrix with large domain, *K*_*L*_, can then by calculated as

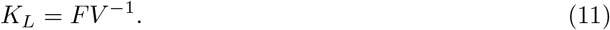

The elements of *K*_*L*_ contain the number of new individuals in each stage and patch produced by one initial new individual in each stages and patch. However, since new individuals are only produced in stage 1, *K*_*L*_ will only have *l* non-zero rows. If equation 10 is arranged so that the populations of stage 1 individuals are in the first *l* rows, then *K*_*L*_ will have a *l* × *l* submatrix in the upper left-hand corner [71]. This submatrix is the next-generation matrix *K*, where the elements of *K, k*_*ij*_, give the number of new (stage 1) individuals produced on patch *i* from one initial stage 1 individual on patch *j*. In equation 10, if the maturation rates from stage *k* to *k* + 1 on patch *j* are 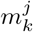, the death rates in stage *k* are *d*^*j*^, the birth rate in the last stage on patch *j* is 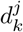, and the probability of successful attachment on patch *i* as a larvae leaving patch *j* is *p*^*ij*^, then the entries of *K* can be written as:

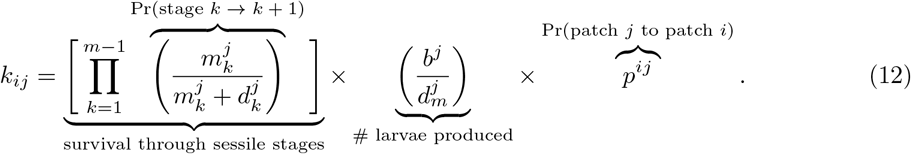

### Example of the next-generation matrix in for a two patch metapopulation

Here we illustrate the construction of the next-generation matrix using a two patch example for a species which has two sessile stages. The population of sessile stage *k* on path *j* is given by 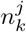 and the maturation rates, death rates, birth rate, probability of dispersal are defined as in the preceding paragraph. The dynamics of the system can be described by

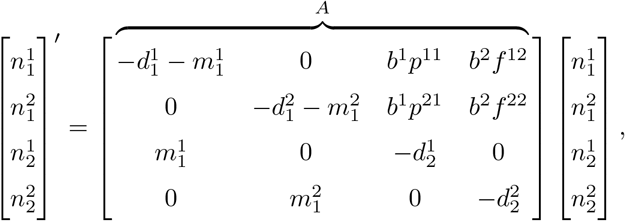

where we have arranged the system of equations so that individuals in the first stage appear in the top rows. We decompose *A* = *F* −*V* where *F* contains all the births of new stage 1 individuals and *V* contains all the remaining transitions, so that *F* and *V* are given by

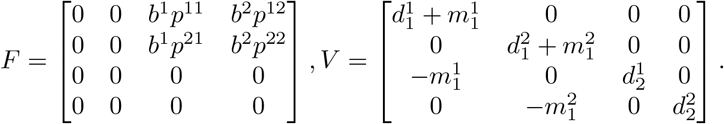

The next-generation matrix with large domain, *K*_*L*_, can then be calculated as *K*_*L*_ = *FV* ^*−*1^, where

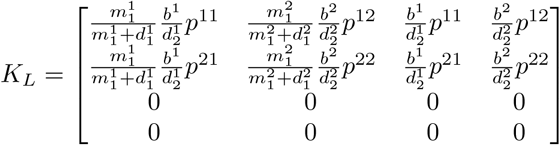

and then the next-generation matrix, *K*, will be given by the 2 × 2 submatrix in the upper left-hand corner of *K*_*L*_, so

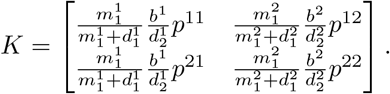

The contribution of patch 1 is therefore

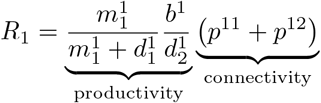

and the contribution of patch 2 is

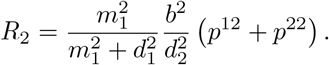

In order for a patch to be a source (*R*_*j*_ *>* 1), both the on-patch productivity and the connectivity to other patches need to multiply to be larger than 1. If either is too low, i.e. if a patch is highly productive but not well connected, or highly connected but not productive, then the patch will be a sink (*R*_*j*_ *<* 1). There are two ways for the entire metapopulation to persist. Either a single patch can persist on its own (*k*_*ii*_ *>* 1), or there must be sufficient production and connectivity within patches such that *R*_0_ = *ρ*(*K*) *>* 1.

## B Calculating the next-generation matrix for discrete time models

The construction of the next-generation matrix for discrete time models is very similar to that of continuous time, though with some minor differences. The population dynamics can now be described by

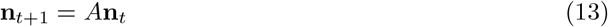

where again **n** is a population vector of size *m* × *l* (number of stages × number of patches) describing the population size on all patches at all sessile stages, but now *A* is a population projection matrix with transition and survival probabilities as well as births.

To calculate the next-generation matrix for the discrete time system, we now decompose *A* = *F* + *T*, where again *F* is a matrix that contains all the birth rates and probabilities of successful of larval dispersal between patches, and now *T* is a matrix that contains all of the survival probabilities and transition probabilities between stages. Under this decomposition, the next-generation matrix with large domain, *K*_*L*_, can by calculated by

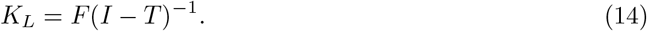

If Equation 13 is arranged so that the populations of all stage 1 individuals are in the top *l* rows, then again the next-generation matrix, *K*, will be the *l* ×*l* submatrix in the upper left hand corner of *K*_*L*_. In the discrete time framework if the probability of transitioning from stage *k* to *k* + 1 is 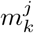, the probability of survival in stage *k* is 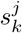, the fecundity in patch *j* is *b*^*j*^, and the probability that a larvae leaving patch *j* successfully arrives on patch *i* is *p*^*ij*^, then the entries of *K* can be written as

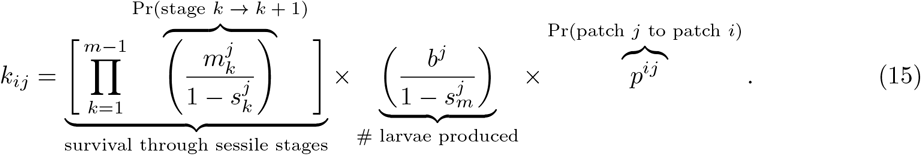

## C Details of the model with age dependent demography in section 2.3

The full set of age density equations for the density of individuals in sessile stage *k* on patch *i* at time *t* and age 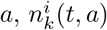, is

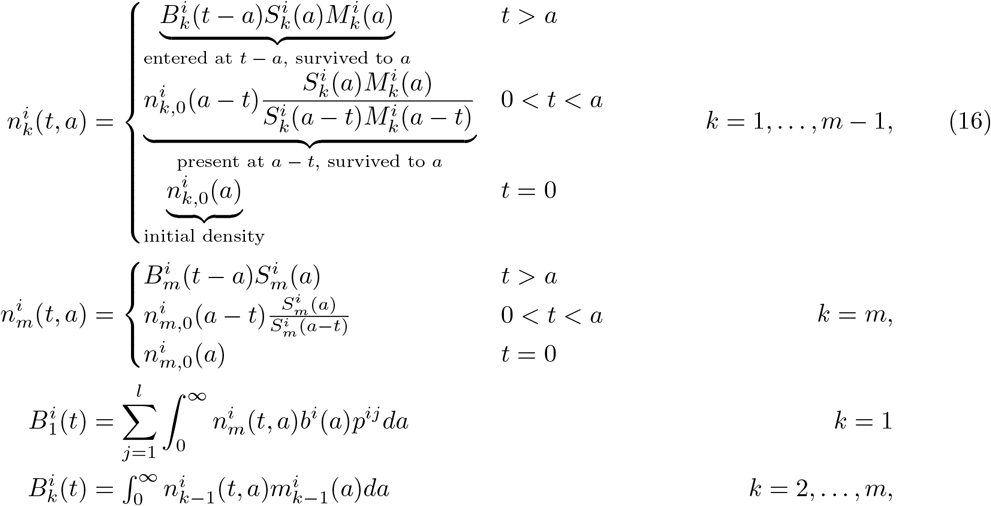

where 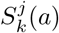 is the probability of survival in stage *k* on patch *j* at stage-age 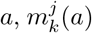 is the maturation rate from stage *k* to *k* + 1, *b*^*j*^(*a*) is the birth rate of larvae on patch *j*, and the probability of a larva leaving patch *j* and successfully attaching on patch *i* is *p*^*ij*^. 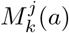 is the probability that an individual has not yet matured from stage *k* to *k* + 1 and can be calculated as 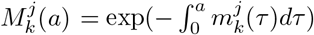. Additional details of the model equations can be found in [23], though there the first larval stage is modelled explicitly and so the set of equations is slightly different.

To construct the next-generation matrix for this system, we need to track the number of stage 1 individuals produced on each patch from one initial stage 1 individual on a given patch. We briefly present the idea of the construction here, and more details can be found in [23].

Let *γ*_*ij*_(*t*) be the rate of production of new stage 1 individuals on patch *i* from an initial stage 1 individual on patch *j*. For an initial individual to be producing offspring it must survive through each of the sessile stages and then produce larvae which settle on another patch. Let *r*_*k*_ be the time that an individual spends in stage *k*. Then in the first *m*− 1 sessile stages *k* = 1, …, *m* − 1 the probability that the individual survives up to *r*_*k*_ is 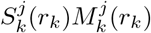 and the rate that they are maturing to the next stage at *r*_*k*_ is 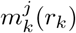. In the last stage the probability that they survive up to *r*_*m*_ is 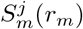 and the rate that they are producing larvae from patch *j* which successfully arrive on patch *i* is *b*^*j*^(*r*_*m*_)*p*^*ij*^.

To calculate the rate of production at time *t, γ*_*ij*_(*t*) we multiply all the survival probabilities, maturation rates, and birth rate in each stage and integrate over all possible *r*_*k*_, where 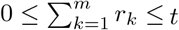. To ensure this bound we rewrite 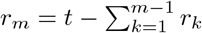. Thus

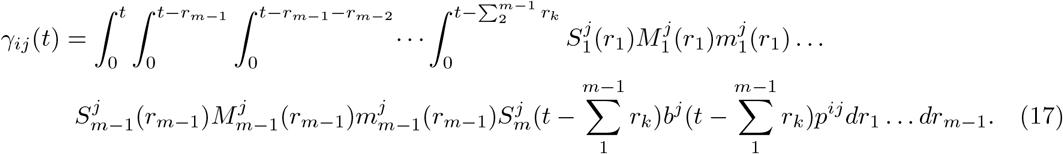

Then we can integrate the rate of production, *γ*_*ij*_(*t*) over all time to find the total number of new individuals produced on *i* from an initial individual on *j*, this is the entry in the *i*th row and *j*th column of the next-generation matrix, *k*_*ij*_. We can use the convolution theorem,

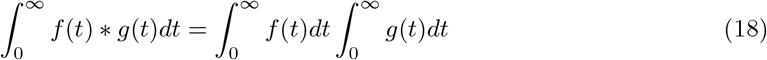

to calculate 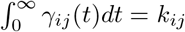 as

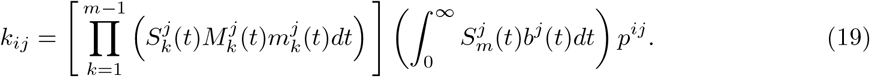

